# Mid-Infrared absorption coefficients of human skin stratum corneum

**DOI:** 10.64898/2026.01.17.699860

**Authors:** Mattia Saita, Alexander Mittelstädt, Nina Köder, Michael Kaluza, Thorsten Lubinski, Dieter Groneberg, Florian K. Groeber-Becker, Werner Mäntele, Sergius Janik

## Abstract

We report infrared absorption coefficients of human stratum corneum from both native *ex vivo* skin and *in vitro* skin models. The spectra show a good transparency in the so-called molecular fingerprint region, even in high relative humidity conditions. Our results provide the quantitative basis for understanding the interaction of Mid-IR light with skin, highly relevant for biomedical sensing and dermatological research.

## 1 INTRODUCTION

The epidermis forms the outermost barrier of human skin, regulating transdermal water loss and protecting against environmental factors such as pathogens, trauma, and tearing. The stratum corneum (SC), as the outermost layer of the epidermis, is the principal component responsible for these barrier properties. A detailed understanding of the SC’s molecular characteristics is crucial for applications such as transdermal drug delivery, cosmetic safety assessment, and the development of non-invasive diagnostic tools, for example the painless transdermal determination of blood or interstitial fluid glucose as an alternative to test strips ^1,2,3,4^.

Infrared spectroscopy offers a powerful method for probing these properties. In the measured spectral range from 4000 to 950 cm^−1^ (approximately 2.5 to 10.5 *µ*m), characteristic absorption signatures can be observed from skin constituents including lipid composition, water content, corneocytes and molecular interactions, both *in vivo* and *ex vivo* ^5,6,7,8,9,10,11^.

Non-invasive infrared spectroscopy of skin has predominantly utilized attenuated total reflectance (ATR) spectroscopy ^7,12^. This technique has been successfully applied to investigate SC hydration ^7,8^ and lipid composition ^13^. However, a significant limitation of the ATR approach is the shallow and ill-defined penetration depth of the evanescent wave, which is typically on the order of a few micrometers and dependent on wavelength, incidence angle, and the refractive indices of the ATR crystal and the sample. Additionally, a perfect contact of the sample to the ATR crystal is strictly required. This renders the determination of quantitative absorption parameters difficult.

In contrast to ATR spectroscopy, transmission spectroscopy provides an optical path length defined only by the sample thickness, enabling the determination of precise absorption coefficients. Such quantitative data are indispensable for the (computational) modeling of light–tissue interactions, which is critical for predicting energy deposition in applications like laser treatments ^4^. Furthermore, accurate optical constants are essential for the design and calibration of novel mid-infrared (Mid-IR) sensors and imaging systems intended for dermatological research and biomedical diagnostics ^14,15,16,17,18,3,19^.

In this letter, we present IR transmission spectra and absorption coefficients of human SC from native ex-vivo samples and human SC obtained from 3D *in-vitro* reconstructed human epidermis (RHE) skin models based on primary cells. The thickness of each stratum corneum sample was estimated via optical coherence tomography (OCT). Finally, we investigated how the absorption coefficient changes at different hydration levels spanning from 20% to 98% relative humidity (RH).

Our results provide a foundational dataset contributing to a deeper understanding of skin-light interaction in the Mid-IR for advancing non-invasive diagnostics, dermatological research, and biomedical sensing applications.

## 2 MATERIALS AND METHODS

### 2.1 3D *in vitro* reconstructed human epidermis models

The RHE models were set up by seeding primary human epidermal keratinocytes onto microporous membrane inserts. Primary human epidermal keratinocytes were isolated from foreskin biopsies from juvenile donors according to the previously published protocol in Ref. ^20^. Cultures were raised to an air-liquid interface (ALI) to promote cellular proliferation and induce terminal differentiation. The models were maintained at the ALI for an extended period of 19–28 days, facilitating the formation of a fully stratified and cornified epidermis of increasing thickness over time. For our subsequent analysis, the mature stratum corneum was mechanically isolated from the RHE samples.

This study was approved by the ethics committee of the University of Würzburg (approval number 182/10 and 280/18) and conducted according to the Helsinki Declaration. Samples were obtained only after the informed consent of the legal guardian(s).

### 2.2 Native human skin explant

Human skin samples from normal adult skin were collected from plastic surgery (abdominoplasty) after informed consent. Prior to mechanical separation of the SC the sample was fixed in a 4% paraformaldehyde solution (ROTI Histofix, Carl Roth GmbH & Co KG, Karlsruhe, Germany; hereafter called Histofix) for 60 min at room temperature. The fixed tissue was subsequently embedded in a medium (Tissue-Tek O.C.T. Compound, Sakura Finetek, USA; herafter called TissueTek) within cryomolds and frozen. Next, the SC was carefully cut down to a thickness of 10 *µ*m with a Cryotome (CM1520, Leica, Nussloch, Germany) and transferred onto a Calcium Fluoride (CaF_2_) window for the Fourier Transform Infrared (FTIR) spectroscopy measurements.

### 2.3 FTIR measurements

FTIR measurements were performed in transmission mode using a compact FTIR spectrometer (Alpha, Bruker Optik GmbH, Ettlingen, Germany). Spectra were acquired by averaging 64 scans at a 4 cm^−1^ spectral resolution over the 4000–950 cm^−1^ range without filters. The skin samples were sandwiched between two wedged CaF_2_ windows (Korth Kristalle GmbH, Altenholz, Germany), using ring-shaped spacers to prevent compression of the sample, see Fig. 1 (a).

**FIGURE 1.**
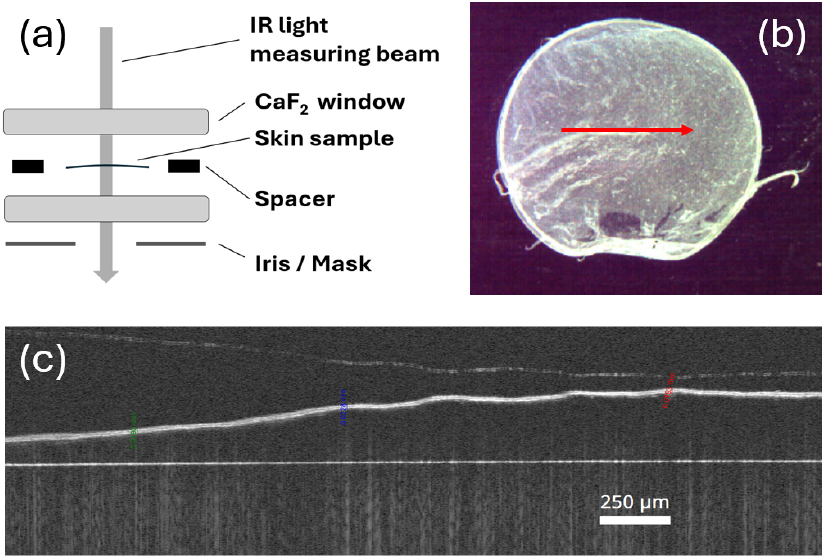
(a) Expanded sketch of the sample sandwich including the description of each component. (b) Image from the OCT-integrated camera of an exemplary RHE sample, the red line indicates where the OCT profile was measured. (c) OCT profile used to measure the skin sample thickness.

For the measurements at different hydration levels, the sample sandwiches were prepared and sealed inside the incubator using a silicon-based sealant (Korasilon, QUAX GmbH, Obernburg, Germany) to prevent leaking of humidity. A background spectrum acquired using empty CaF_2_ windows was subtracted from each sample spectrum. Custom-built optical masks were used to prevent any false light from reaching the detector. Measurements of thicker RHE samples (> 25 *µ*m), where peak absorbance exceeded 3 OD, were excluded from the analysis, as excessive absorption can increase noise and distort spectral band shapes. A linear baseline correction was applied using OPUS software’s (Bruker Optik GmbH, Obernburg, Germany) built-in function to compensate for scattering artifacts. For this correction, we assumed that the sample’s absorption at 4000 cm^−1^ and 1850 cm^−1^ is zero.

### 2.4 Incubation and estimation of relative humidity

Absorption spectra of five distinct RHE SC samples were investigated at various RH levels. Briefly, samples were placed on CaF_2_ windows and incubated in a CO_2_ incubator (BBD6220, Life Technologies GmbH, Darmstadt, Germany), used solely as hydration chamber, for 45 minutes at several RH levels ranging from 20–98 % at 30 ° C. The samples were then sealed with the second window inside the incubator and measured in-situ; see Sec. 2.3. Care was taken to ensure the sample humidity, measured with a thermohygrometer (625, Testo SE & Co. KGaA, Freiburg, Germany), would not change during the sealing process and the RH reported was recorded from the sensor at the moment of sealing. While this procedure introduces minor uncertainty in absolute RH, this experiment focuses on broad hydration changes, rendering those variations negligible for the present analysis.

The 45-minute incubation time was chosen after a comparative analysis with a 12-hour period yielded equivalent results. Potential humidity leaks during measurements were ruled out, as repeated measurements over time demonstrated consistent spectral reproducibility; see Supporting Information.

### 2.5 OCT

OCT imaging was performed using a Thorlabs Ganymede Series system, OCTG9 scanner, (THORLABS GmbH, Bergkirchen, Germany) with a central wavelength of 880 nm and an axial resolution of 3 *µ*m in water. For the RHE samples and native human skin explants, measurements were acquired at a minimum of three distinct sites along a linear path. The mean value from these sites was used for subsequent analysis; see Fig. 1 (b, c). To calculate the thickness of the stratum corneum, a refractive index of *n* = 1.54 was applied ^21,22,23,24^.

## 3 RESULTS

### 3.1 Comparison of native skin and in vitro RHE skin models

Native human SC was measured in transmission to determine its Mid-IR absorption coefficient. Practical and ethical considerations restrict the availability of native human skin samples. Additionally, native skin samples show a variability (e.g., donor patient age, body location of surgical site) and require fixation with additives; see Sec. 2.2. In contrast, RHE models offer a greater uniformity and can be readily prepared in the lab without exogenous additives.

Here, we demonstrate the validity of RHE SC as a surrogate for native human SC for determining its Mid-IR absorption coefficient. For clarity and consistency, the results from a single, representative RHE sample are presented in Figs. 2 and 4, as all RHE samples yielded equivalent spectra, within the measurement error.

**FIGURE 2.**
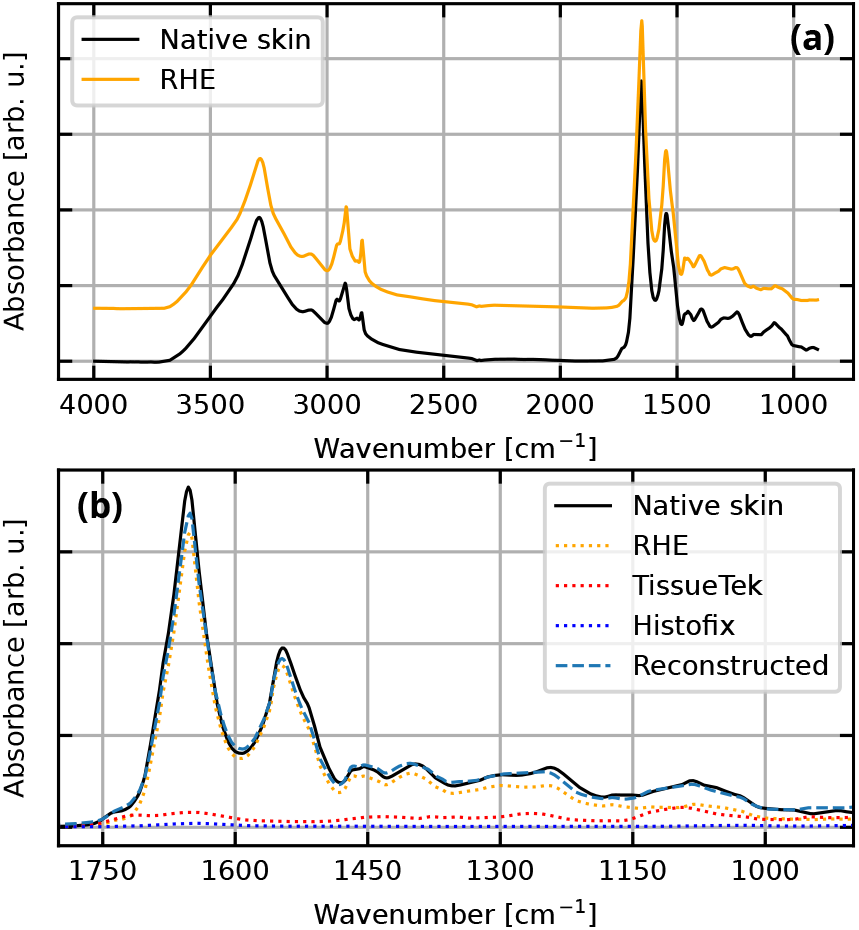
(a) The spectra of native skin and RHE are compared. The RHE spectrum has been offset to improve visualization. (b) A least-squares fit of the native skin spectrum in the range of 1800–900 cm^−1^ as a linear combination of the RHE spectrum and the spectra of the additive solutions.

In Fig. 2 (a), the absorption spectra are compared. The overall spectral shape is well preserved in the RHE sample, with minor differences in the Amide I (1700–1600 cm^−1^) and fingerprint (1500–950 cm^−1^) regions. As shown in Fig. 2 (b), these discrepancies are attributed to additives required for native skin preparation. Thus, the native skin spectrum is accurately described by a spectral combination of the RHE spectrum and the two additives TissueTek and Histofix; see Sec. 2.2).

The fit demonstrates that the broader Amide I band in the native skin spectrum is due to both additives showing broad bands in the same range. Similarly, the two broad absorption features of the native skin around 1240 cm^−1^ and 1080 cm^−1^ are well described by the superposition of the RHE absorption and the contribution of the additives. The precise chemical composition of the proprietary products TissueTek and Histofix is not available, but their broad absorption bands at 1100–1030 cm^−1^, 1260 cm^−1^, 1430 cm^−1^ and 1714 cm^−1^ are compatible with CO stretching, O-H bending, and C=O stretching vibrations of alcohols, carbonyl or ester groups.

This comparison shows that using the native skin spectrum to determine its absorption coefficient would be inaccurate due to the presence of additives. Furthermore, it demonstrates that RHE models provide a reliable and accurate representation of human SC absorption and can be confidently used to determine optical properties of human skin SC.

### 3.2 Absorption coefficients of dry human stratum corneum

We recorded infrared absorption spectra of eight different RHE SC samples from two different batches and two different preparations (see Sec. 2). The spectra were measured at room humidity (<55 RH% for all samples). The sample thicknesses ranged from 12–24 *µ*m, as estimated with OCT; see Sec. 2.5. Fig. 3 shows the calculated absorption coefficients for each sample.

**FIGURE 3.**
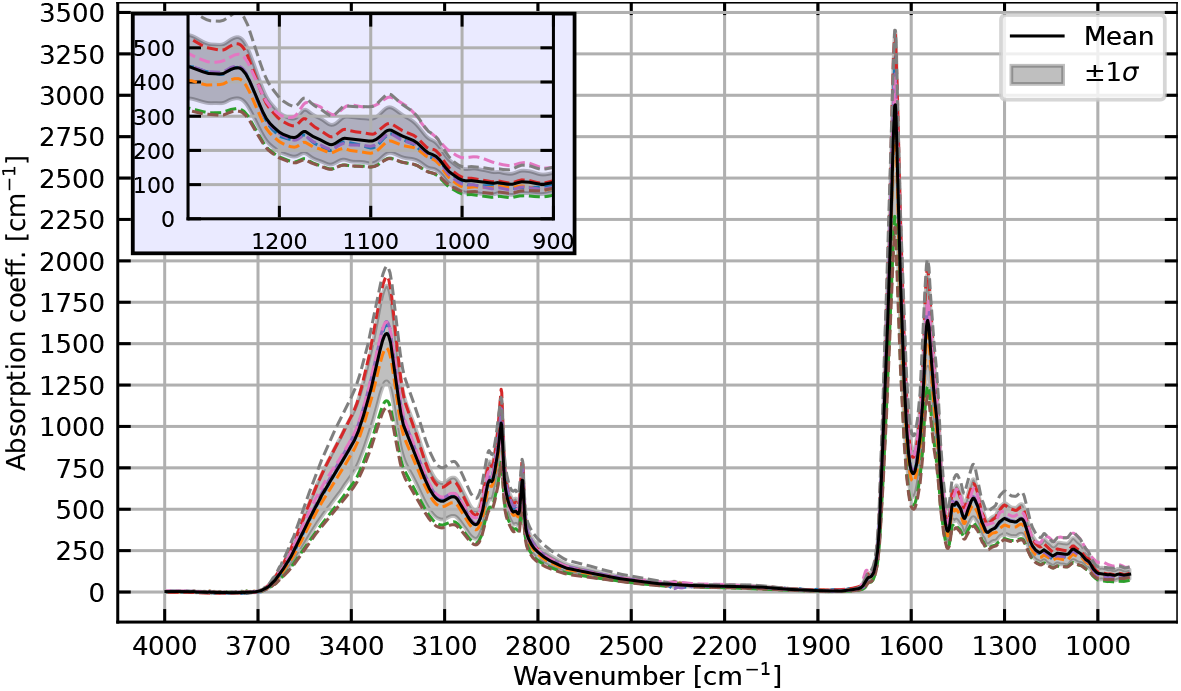
Absorption coefficients of all RHE stratum corneum samples measured (dashed) as well as the mean (black) and corresponding 1*σ* band (gray shaded). The inset shows a zoom into the spectral region of 1300–900 cm^−1^.

**FIGURE 4.**
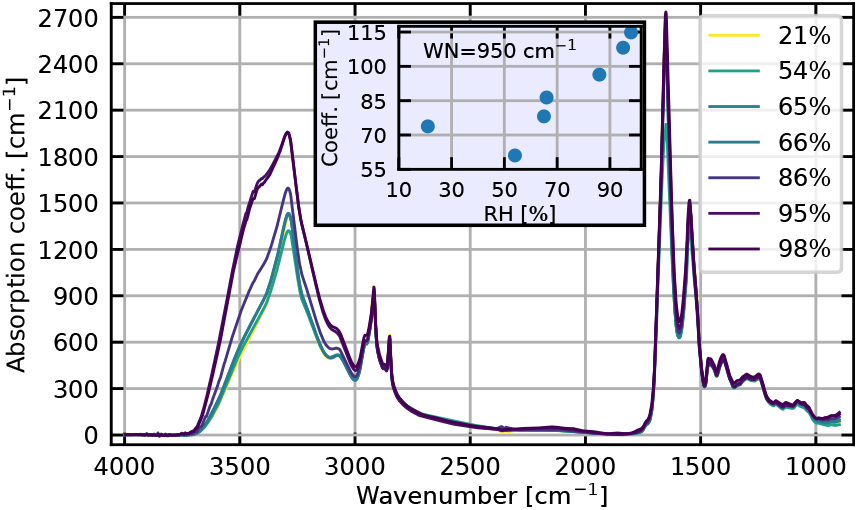
Absorption coefficients at different levels of RH. The inset shows the absorption coefficients at wavenumber = 950 cm^−1^ versus the RH.

The slight variations in the calculated absorption coefficients are attributed to uncertainties in thickness estimation, sample preparation, and scattering corrections. Computing the mean and standard deviation enhances the robustness of the investigation and provides a confidence interval for the full spectral range.

Our results are in good agreement with literature reports of human SC measured in FTIR microspectroscopy and with human keratin spectra ^25^. The values we report are in line with the previously estimated absorption coefficient in Ref. ^16^ in the spectral region 1020–1100 cm^−1^ based on the water content of SC.

The absorption coefficients reported here, 150–250 cm^−1^ for the same range, are derived from well-defined thickness measurements and hydration states, thus providing a more accurate determination.

SC components, such as keratin filaments, may exhibit a preferred orientation, potentially introducing anisotropy ^26^. The absorption coefficients reported here were obtained using an unpolarized beam with the electric field perpendicular to the propagation direction, as is relevant for typical applications of incident light on skin.

### 3.3 Impact of relative humidity on the spectra

Fig. 4 shows the absorption coefficients for five different RHE SC samples at various hydration levels. As expected, the spectra exhibit the most significant changes around 3300 cm^−1^, the strong water OH stretching vibration band, which is in good agreement with a report on keratin hydration in Ref. ^27^. A smaller hydration effect is observable below 980 cm^−1^, the water libration band. In general, hydration levels from 20–98 RH% do not significantly impact the absorption coefficients in the 3000–980 cm^−1^ spectral region. Although a change was also expected near 1650 cm^−1^, the water OH bending vibration, this region is dominated by the prominent keratin Amide I band, potentially obscuring water contributions.

The inset shows the absorption coefficient at 950 cm^−1^ as a function of hydration; data for other wavenumbers are available in the Supporting Information. Despite minor single-point deviations, the data show a clear trend of water absorption by the SC sample exposed to different RH levels consistent with early literature reports ^28,29^.

## 4 CONCLUSION

We presented a determination of the absorption coefficient of human skin SC in the 4000 to 950 cm^−1^ (approximately 2.5 to 10.5 *µ*m) spectral range and investigated its dependence on hydration. The SC’s Mid-IR absorption is dominated by strong protein bands from its high concentration of keratin filaments. Our data shows that the absorption coefficient in the 3000– 980 cm^−1^ spectral range remains minimally affected by the RH, even at humidity levels approaching 98%, underscoring the SC’s function as a hydrophobic barrier.

The absorption coefficient of the SC is the critical parameter that dictates light penetration depth and light-skin interaction for Mid-IR-based biomedical sensing applications and dermatological research. Our findings provide a fundamental data set necessary to quantify this light-skin interaction accurately. In view of the application of mid-infrared sensor techniques for transdermal measurements of metabolic parameters, e.g. skin glucose, the measurements presented here clearly indicate that a mid-infrared measuring beam in the wavenumber range 950– 1200 cm^−1^ penetrates deep enough into skin layers to probe interstitial fluid glucose, given the low absorption coefficient of SC in that range.

This contribution will directly benefit medical diagnostics, therapy and cosmetics, enabling the design and validation of novel Mid-IR-based sensors and revolutionizing wearable health technology for continuous, non-invasive monitoring of biomarkers like glucose or skin health.

## Abbreviations

SC: stratum corneum
OCT: optical coherence tomography
ATR: attenuated total reflectance
FTIR: Fourier transform infrared
RHE: in vitro reconstructed human epidermis
RH: Relative Humidity
ALI: air-liquid interface

## AUTHOR CONTRIBUTIONS

M.S. and A.M. contrubuted equally to this work and performed all the measurements, the data analysis and wrote the manuscript with input from all authors. S.J. conceived the idea, initiated the collaboration with ISC and supervised the project. WM helped writing the manuscript and provided scientific supervision. N.K. prepared the skin samples and helped with the measurements at Fraunhofer ISC. D.G. was involved in the planning and supervised the work at Fraunhofer ISC. T.L., M.K and F.K.GB. were in charge of overall direction and planning.

## ACKNOWLEDGMENTS

The authors would like to thank Christa Albert and Annika Baumann who supported the sample preparation as well as Honorata M. Ropiak and Peter Lachmann for microscopy images of the skin samples.

## ETHICS APPROVAL STATEMENT

The use of primary human cells for the reconstruction of the RHE was approved by the ethics committee of the university of Würzburg under protocol number (approval number 2018-280_5-dvh), in accordance with the declaration of Helsinki and institutional ethical guidelines.

## PATIENT CONSENT STATEMENT

Donor skin samples were obtained after written informed consent for the use of the material in research.

## SUPPORTING INFORMATION

Additional supporting information may be found in the online version of the article at the publisher’s website.

## APPENDIX

